# The draft nuclear genome assembly of *Eucalyptus pauciflora*: new approaches to comparing *de novo* assemblies

**DOI:** 10.1101/678730

**Authors:** Weiwen Wang, Ashutosh Das, David Kainer, Miriam Schalamun, Alejandro Morales-Suarez, Benjamin Schwessinger, Robert Lanfear

**Author notes:** Equal contribution (Weiwen Wang, Ashutosh Das). **Email**: Weiwen Wang, Ashutosh Das, David Kainer, Miriam Schalamun, Alejandro Morales-Suarez, Benjamin Schwessinger, Robert Lanfear.

## Abstract

**Background:** Selecting the best genome assembly from a collection of draft assemblies for the same species remains a difficult task. Here, we combine new and existing approaches to help to address this, using the non-model plant *Eucalyptus pauciflora* (snow gum) as a test case. *Eucalyptus pauciflora* is a long-lived tree with high economic and ecological importance. Currently, little genomic information for *Eucalyptus pauciflora* is available.

**Findings:** We generated high coverage of long-(Nanopore, 174x) and short-(Illumina, 228x) read data from a single *Eucalyptus pauciflora* individual and compared assemblies from four assemblers with a variety of settings: Canu, Flye, Marvel, and MaSuRCA. A key component of our approach is to keep a randomly selected collection of ~10% of both long- and short-reads separate from the assemblies to use as a validation set with which to assess the assemblies. Using this validation set along with a range of existing tools, we compared the assemblies in eight ways: contig N50, BUSCO scores, LAI scores, assembly ploidy, base-level error rate, computing genome assembly likelihoods, structural variation and genome sequence similarity. Our result showed that MaSuRCA generated the best assembly, which is 594.87 Mb in size, with a contig N50 of 3.23 Mb, and an estimated error rate of ~0.006 errors per base.

**Conclusions:** We report a draft genome of *Eucalyptus pauciflora*, which will be a valuable resource for further genomic studies of eucalypts. These approaches for assessing and comparing genomes should help in assessing and choosing among many potential genome assemblies for a single species.

## Data Description

### Introduction

Eucalypts are widely distributed in Australia, including three genera *Eucalyptus, Corymbia* and *Angophora*, and have around 900 species [1]. *Eucalyptus pauciflora* (*E. pauciflora*) (Fig. 1), also known as snow gum, is a highly variable eucalyptus species that inhabits diverse landscapes in south-eastern Australia [1]. *E. pauciflora* can survive from close to sea level to up to the tree line of the Australian Alps, displaying the broadest altitudinal range in the *Eucalyptus* genera [2–4]. Due to its wide distribution and drought and cold tolerance, *E. pauciflora* is used for carbon offset plantings, ecological restoration, honeybee food source, and also has medicinal uses [1, 5–11]. However, genomic resources for *E. pauciflora* are currently very limited: there exists a single chloroplast genome [12], two sets of microsatellite markers [13, 14], and two nuclear loci used for phylogenetics [15]. The assembly of *E. pauciflora* genome will assist in elucidating the genetic basis of cold tolerance in *Eucalyptus*.

**Figure 1:**
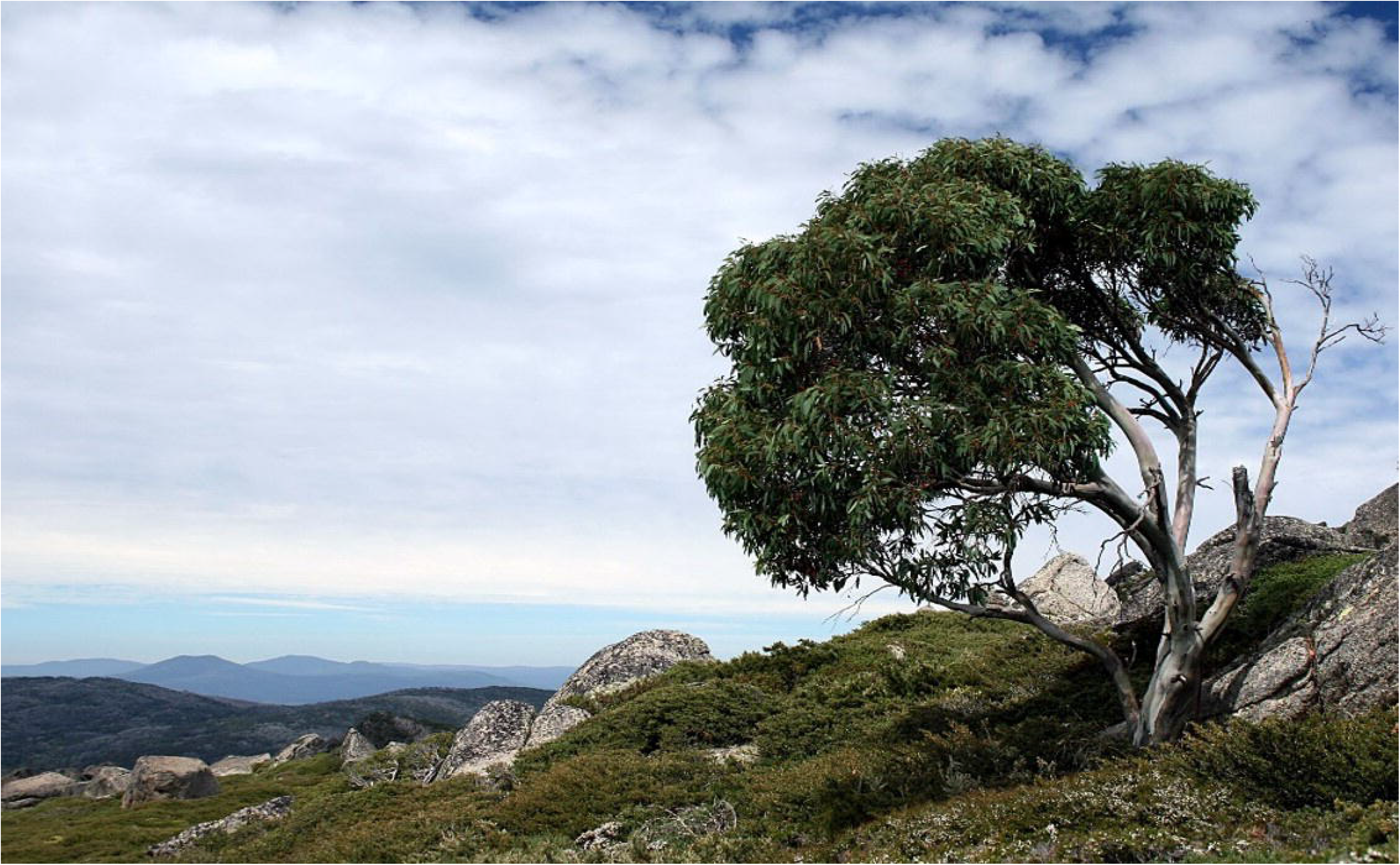
The *E. pauciflora* sequenced in this study. This *E. pauciflora* is located in Thredbo, Kosciuszko National Park, New South Wales, Australia (36° 29’ 39.58” N, 148° 16’ 58.73’ E).

Across the ~900 extant eucalypt species, there are only two genomes published: those for *E. grandis* and *E. camaldulensis* [16, 17]. Both of these genomes were sequenced with a combination of Sanger sequencing and short-read sequencing, and as a result both assemblies are somewhat fragmented. There are 81,246 scaffolds in *E. camaldulensis* assembly [17]. While the *E. grandis* genome is highly contigous, assembled to chromosom e level, it still has 4,941 unplaced scaffolds [16]. New technologies, such as third-generation long-read sequencing, have the potential to produce less fragmented assemblies at a fraction of the cost of previous methods. Nevertheless, many challenges still remain, not least of which is that different genome assembly software, and small changes to the parameters of a single piece of software, can produce substantially different assemblies. In light of this, methods for choosing the most accurate assembly from a set of possible assemblies have become increasingly important.

Two metrics are commonly used to assess and compare genome assemblies: contig N50 and Benchmarking Universal Single-Copy Orthologs (BUSCO [18], RRID:SCR_015008) scores. The contig N50 is the size of the contig where at least 50% of the assembled nucleotides can be found in contigs of that size or larger. The N50 is a measure of genome contiguity, where a higher N50 suggests a genome that has been assembled into fewer and larger contigs. All else being equal, we should prefer genome assemblies with a larger N50, up to the point where the N50 is equal to the N50 of the chromosomes themselves. Perhaps because of this, the N50 is one of the most widely reported metrics in genome assembly. However, it is important to remember that the N50 measures contiguity, not accuracy. For example, N50 scores may be artificially inflated by incorrectly linking contigs [19, 20]. The BUSCO score estimates the proportion of highly conserved orthologous genes that are present in assemblies. The underlying assumption is that there exists a certain set of highly conserved single-copy genes, the vast majority of which we should expect to observe in single copies in any given haploid genome assembly. BUSCO scores provide a very useful measure of genome assembly completeness (a component of accuracy), and in principle we should prefer genome assemblies with BUSCO scores closer to 100%. One limitation of BUSCO scores is that they assess only a very small proportion of the genome, typically around 1000 highly conserved genes which represent less than 1% of the total genome. Furthermore, by their nature these protein-coding regions of the genome tend to be among the easiest to assemble because they are usually single-copy regions of high complexity. Hence, assemblies can have very similar BUSCO scores even if they differ considerably in their assembly of the non-BUSCO genomic regions, which means that it is sometimes difficult to use BUSCO scores to distinguish among competing assemblies [21]. In this study, we complement these commonly-used measures with a range of other metrics to assess and compare genome assemblies, and we use these measures to choose the best draft assembly of *E. pauciflora*.

One measure we propose is the assembly ploidy: the proportion of the genome that is represented by haploid contigs. One important problem in genome assembly is that we commonly represent the genome of diploid (or polyploid) organisms as a haploid sequence. Traditionally, genome projects would alleviate this problem by sequencing highly inbred individuals [22, 23], thus reducing the discrepancy between the diploid individual and the haploid representation. However, as genome assembly has become more commonplace, we often want to assemble the genomes of highly heterozygous individuals. For example, heterozygosity in *Eucalyptus* is around 1% [24], and varies substantially along the genome [16]. The consequence of this is that regions of low heterozygosity tend to be assembled into a single collapsed haploid sequence, whereas regions of high heterozygosity tend to be assembled into two haplotypes of the same region, which are usually labelled the ‘primary contig’ (referring to the longer of the two contigs) and the ‘haplotig’ (referring to the shorter of the two contigs) [25]. Although there has been some progresses in estimating truly diploid assemblies [25, 26], most assemblers still produce primary contigs and haplotigs without labelling them as such [27, 28]. Crucially, unidentified haplotigs may cause issues in the downstream analyses, because many analyses assume that we have a haploid representation of the genome. Because of this, we propose a novel and simple (but imperfect) metric to measure the assembly ploidy, which is simply the ratio of the assembly size to the estimated haploid genome size. If the aim is to produce a haploid representation of a genome, then an assembly ploidy of 1 is preferable (i.e. the assembly size should equal the estimated haploid genome size). If the aim is to produce a diploid representation of a genome, then an assembly ploidy of 2 is preferable (i.e. the assembly size should be double the estimated haploid genome size). One limitation of this metric is that it is sensitive to errors in the estimation of haploid genome size, and it is also sensitive to errors in genome assembly (e.g. highly incomplete assemblies) that might affect the numerator. Nevertheless, in combination with other measures, we show below that the assembly ploidy provides a useful metric with which to compare genome assemblies.

We also apply a suite of measures designed to provide a genome-wide assessment of contiguity and accuracy that can complement the widely-used contig N50 and BUSCO scores. The advantages of these measures lie in the fact that they assess more of the genome than BUSCO scores, though each also has its limitations. The first measure is the long-terminal repeat (LTR) assembly index, or LAI [21]. The LAI score is the proportion LTR sequences in the genome that are intact, and is independent of genome size and repeat content. In general, a higher LAI score suggests a more contiguous and complete assembly [21]. The second measure we use is the base-level error rate evaluated by remapping independent sets of long and short validation reads (around 10% of all reads, randomly selected) to the assembly. Previous studies have evaluated the base-level error rate by remapping all reads to the assembly [29, 30]. Here, we use validation reads which are not involved in the assembly, in order to avoid any possible biases introduced by validating an assembly with the same data that was used to produce it. For a perfect assembly in which the ploidy of the entire assembly matches the ploidy of the individual, a lower base-level error rate is preferable, with a theoretical minimum of the error rate of the sequencing technology (e.g. ~0.3% for raw Illumina reads [31], and ~10-15% for raw Nanopore reads [32, 33]). For a haploid representation of a diploid assembly, the minimum possible base-level error rate will be higher, because by necessity a haploid representation of a heterozygous site will not match approximately half of the reads. In this case, the theoretical minimum base-level error rate is the sum of the error rate of the sequencing technology and half of the heterozygosity. The third measure is the computing genome assembly likelihoods (CGAL) score [20]. The CGAL score is the likelihood of an assembly calculated from a model that accounts for errors in reads, read coverage across the assembly, and the proportion of reads that do not contribute to the assembly. A higher likelihood suggests that a genome assembly is a better representation of the truth. The fourth measure we use is the number of structural variants detected when re-mapping our long validation reads to assemblies. As with the base-level error rate, if the ploidy of the assembly matches the ploidy of the individual, then the theoretical minimum of this metric is the structural error rate introduced into sequencing reads by the sequencing technology. For a haploid representation of a diploid genome, the theoretical minimum is the sum of the error rate of the technology plus half of the structural heterozygosity. These two quantities are rarely known, but nevertheless, a very high structural error rate of validation reads mapped to a haploid assembly may indicate cases in which the assembly has a large proportion of incorrectly linked contigs. The final measure is the genome sequence similarity of each assembly when compared to all other assemblies. This measure does not provide any information relative to an underlying truth, but it may help to identify significant differences between otherwise plausible genome assemblies that can aid in choosing the best assembly. The selection of the best assembly should consider all measures together.

Here, we used long- and short-reads to create a draft haploid assembly of the *E. pauciflora* genome. We use the metrics we describe above to compare a range of assemblies from a range of different assemblers. We performed different assemblies with long-read-only assemblers (Canu (Canu, RRID:SCR_015880) [34], Flye [35] and Marvel [36]) and hybrid assembler MaSuRCA (MaSuRCA, RRID:SCR_010691) [37], using long-read datasets with different minimum read lengths in each case (1 kb and 35 kb).

### Sample collection, DNA sequencing and quality control

We collected leaves from the single *E. pauciflora* tree near Thredbo, Kosciuszko National Park, New South Wales, Australia (36° 29’ 39.58” N, 148° 16’ 58.73” E) in March 2016 (for Illumina sequencing) and June 2017 (for MinION sequencing). We stored leaves at 4°C when transported them to the laboratory.

For long-read sequencing, we extracted high molecular weight genomic DNA from leaves following a protocol optimized for *Eucalyptus* nanopore sequencing [38]. We prepared ONT 1D ligation libraries according to the manufacturer’s protocol (SQK-LSK108) and sequenced the reads using MinKNOW v1.7.3 with R9.5 flowcells on a MinION sequencer. We performed basecalling with Albacore v.2.0.2 (Albacore, RRID:SCR_015897). This resulted in 12,584,100 raw long-reads (106.96 Gb) with average read length of 8.5 kb. We removed adapters from long-reads with Porechop v0.2.1 (Porechop, RRID: SCR_016967) [39]. Next, we trimmed bases with quality <10 on both ends of the reads using NanoFilt (NanoFilt, RRID:SCR_016966) [40] and discarded reads shorter than 1 kb after trimming. This recovered 96.66 Gb of long-read data comprising 7,711,141 filtered reads with an average read length of 12.53 kb (minimum 1 kb and maximum ~150 kb). Given an estimated genome size of 500 Mb (see below), this represents a coverage of 193x.

For short-read sequencing, we extracted genomic DNA from freeze-dried leaves using a CTAB protocol [41] followed by purification with a Zymo kit (Zymo Research Corp). We constructed TruSeq Nano libraries with an insert size of 400 bp using protocol provided by Illumina, then sequenced the reads (paired-end 150 bp) using an Illumina Hiseq2500 platform (Illumina Inc., San Diego, CA). This Illumina sequencing generated 506,840,789 paired raw reads (152.05 Gb). We used BBDuk v37.31 (BBmap, RRID:SCR_016965) [42] to remove adapters and to trim both sides of raw short-reads which quality was lower than 30. We discarded filtered reads with a length under 50 bp. Around 122.69 Gb short-read data containing 414,697,585 paired reads were left, representing 246x coverage with an estimated genome size of 500 Mb (see below).

### Genome size and heterozygosity estimation

We used GenomeScope (GenomeScope, RRID:SCR_017014) [43] and SGA-preqc (SGA, RRID:SCR_001982) [44] to estimate the *E. pauciflora* genome size. We first generated a 32-mer distribution using Jellyfish v1.1.12 (Jellyfish, RRID:SCR_005491) [45] from all of our short-reads, then ran GenomeScope using this 32-mer distribution with a maximum k-mer coverage of 1000x. This gave a genome size estimate of 408.16 Mb (Additional file 1: Fig. S1), which is lower than expected for other *Eucalyptus* species [16, 17]. However, it is known that genomic repeats can lead to underestimation of genome sizes from uncorrected kmer distributions [46], and the *Eucalyptus* genome is repeat-rich, for example around 50% of genome was annotated as repeats in *E. grandis* [16], suggesting that 408.16 Mb may be a significant underestimate of the genome size. SGA-preqc estimates genome size from k-mer distributions that are corrected to attempt to better account for repeat content, in line with this, SGA-preqc gave a genome size estimate of 529.40 Mb. Because of this, we expect that the SGA-preqc genome size is likely to be more accurate, and in what follows we assume that the *E. pauciflora* genome size is roughly 500 Mb. This suggests that the *E. pauciflora* genome may be around ~30% smaller than that of the other two sequenced *Eucalyptus* species, *E. grandis* (691.43 Mb) [16] and *E. camaldulensis* (654.92 Mb) [17]. However, the genome sizes of *E. grandis* and *E. camaldulensis* may be overestimated due to the assembly and scaffolding of both haplotypes at heterozygous regions.

### Creation of assembly and validation datasets

We separated our long-read and short-read data into assembly dataset (~90% of reads) and validation dataset (~10% of reads) by randomly assigning the trimmed and filtered reads into the two datasets. The assembly dataset comprised 86.94 Gb of long-read data (174x coverage) and 114.10 Gb of short-read data (228x coverage). The validation dataset comprised and 9.67 Gb of long-read data (19x coverage) and 8.59 Gb of short-read data (17x coverage).

### Genome assembly

Here, we compared five long-read-only assemblies and two hybrid assemblies. For each combination of data and genome assembler, we followed the same genome assembly pipeline. We first used the assembler to produce an initial assembly. Following this, we identified and removed contigs from contaminant sequences, and then polished the resulting assembly. We then identified and removed haplotigs from the assembly, and finally re-polished each assembly after haplotig removal. To select the best assembly, we calculated the contig N50 with Quast [19], BUSCO scores with BUSCO, and LAI scores using the LTR_retriever pipeline [47]. After mapping the long- and short-validation reads to the final assemblies (using Ngmlr [48] for the former and Bowtie2 (Bowtie2, RRID:SCR_016368) [49] for the latter), we calculated the base-level error rate using Qualimap [50] the structural variant error rate using Sniffles [48], and CGAL scores using CGAL. Finally, we performed whole genome alignment between different assemblies with NUCmer module of MUMmer [51].

Oxford Nanopore reads tend to have error rates of ~10-15%, which can make assembly of uncorrected reads very challenging. To alleviate this, we first corrected the long-reads assembly dataset with Canu v1.6 with default parameters except for setting corMinCoverage to 8, meaning that read correction would only be applied where at least 8 reads overlapped. We deemed this reasonable given the very high coverage of our data (174x). We then put the corrected long-read datasets into two sets for assembly. The first dataset contained all corrected long-reads, such that the minimum read length was 1 kb (174x of coverage). The second dataset contained all corrected reads longer than 35 kb (~40x of coverage). We refer to these datasets as the 1 kb and the 35 kb datasets, respectively.

We attempted six long-read-only assemblies and two hybrid assemblies. Assemblies solely with long-read data were performed on corrected reads of two read lengths (1 kb and 35 kb) using three long-read assemblers: Canu v1.6 and v1.7, Flye v2.3.5 and Marvel v1.0. The Marvel assembly with 1kb dataset was not feasible because it required more disk space than we had available, resulting in five successful long-read only assemblies. We used MaSuRCA v3.2.6 to perform hybrid assemblies with both read length datasets (1 kb and 35 kb) each combined with the short-read dataset. In what follows, we refer to these assemblies as Canu_1kb, Canu_35kb, Flye_1kb, Flye_35kb, Marvel_35kb, MaSuRCA_1kb and MaSuRCA_35kb. We used default settings in all assemblers, and an estimated genome size of 500 Mb where this setting was required. For Canu assemblies, the 1 kb dataset was assembled using Canu v1.6, whereas the 35 kb dataset was assembled using Canu v1.7. We did not repeat the Canu_1kb assembly after Canu v1.7 was released, because we no longer had sufficient computational resources. The chloroplast genome and mitochondrial genome were removed from each assembly. For each assembly, we recorded the runtime in CPU hours, the raw assembly length, and the N50 (Table 1).

**Table 1.**
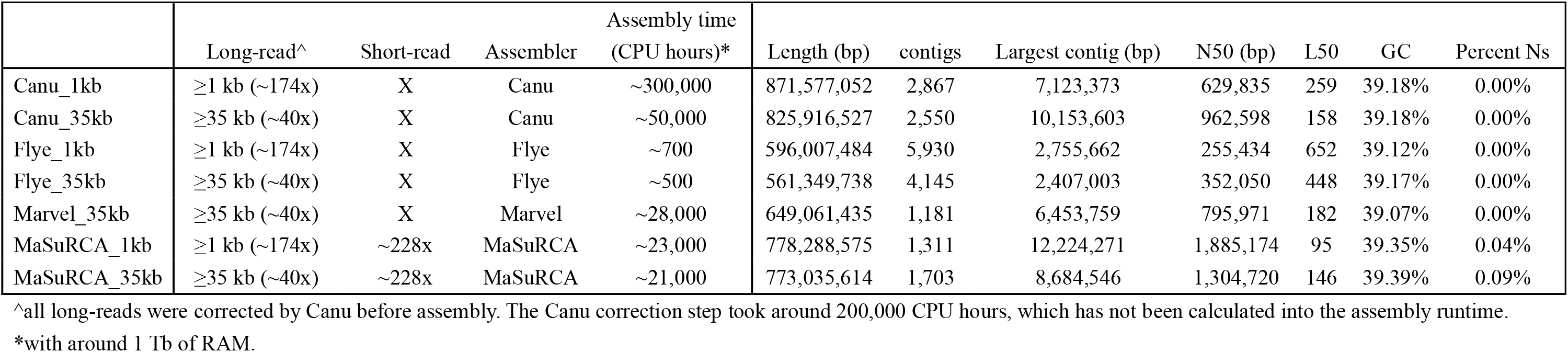
The statistics information of raw assemblies.

|  | Long-read^ | Short-read | Assembler | Assembly time<br>(CPU hours)* | Length (bp) | contigs | Largest contig (bp) | N50 (bp) | L50 | GC | Percent Ns |
| --- | --- | --- | --- | --- | --- | --- | --- | --- | --- | --- | --- |
| Canu_1kb | ≥1 kb (~174x) | X | Canu | ~300,000 | 871,577,052 | 2,867 | 7,123,373 | 629,835 | 259 | 39.18% | 0.00% |
| Canu_35kb | ≥35 kb (~40x) | X | Canu | ~50,000 | 825,916,527 | 2,550 | 10,153,603 | 962,598 | 158 | 39.18% | 0.00% |
| Flye_1kb | ≥1 kb (~174x) | X | Flye | ~700 | 596,007,484 | 5,930 | 2,755,662 | 255,434 | 652 | 39.12% | 0.00% |
| Flye_35kb | ≥35 kb (~40x) | X | Flye | ~500 | 561,349,738 | 4,145 | 2,407,003 | 352,050 | 448 | 39.17% | 0.00% |
| Marvel_35kb | ≥35 kb (~40x) | X | Marvel | ~28,000 | 649,061,435 | 1,181 | 6,453,759 | 795,971 | 182 | 39.07% | 0.00% |
| MaSuRCA_1kb | ≥1 kb (~174x) | ~228x | MaSuRCA | ~23,000 | 778,288,575 | 1,311 | 12,224,271 | 1,885,174 | 95 | 39.35% | 0.04% |
| MaSuRCA_35kb | ≥35 kb (~40x) | ~228x | MaSuRCA | ~21,000 | 773,035,614 | 1,703 | 8,684,546 | 1,304,720 | 146 | 39.39% | 0.09% |

^all long-reads were corrected by Canu before assembly. The Canu correction step took around 200,000 CPU hours, which has not been calculated into the assembly runtime.
\*with around 1 Tb of RAM.

### Contamination detection

Following initial assembly, we used Blobtools [52] to assess contamination in each genome assembly. To do this, we first generated a hit file for each assembly by searching all contigs against the National Center for Biotechnology Information (NCBI) non-redundant nucleotide database using BLASTN v2.7.1+ (BLASTN, RRID:SCR_001598) [53] (E-value ≤ 1e-20). We then analysed the hit file for each assembly using Blobtools, which provides taxonomic annotations and other diagnostic plots to detect contamination in raw genome assemblies. The top-hit was streptophyta phylum, comprising 99.72% to 100% of the hits in different assemblies (Additional file 2: Fig. S2), indicating that there was no potential contamination from a non-plant origin in each raw assembly.

### Genome polishing

We polished each initial genome assembly in order to improve its accuracy. For the Canu, Flye, and Marvel assemblies (i.e. those built from long-reads only), we polished first with Racon [54] using Ngmlr v0.2.6 using the long-read assembly dataset, and then with Pilon v1.22 (Pilon, RRID:SCR_014731) [55] using Bowtie v2.3.4.1 with the short-read assembly dataset. For the MaSuRCA assemblies, we polished only with Pilon because MaSuRCA is a hybrid assembler, and using error-prone long-reads to polish hybrid assemblies tends to induce more errors rather than remove them (Additional file 3: Table S1).

We ran each polishing algorithm for multiple iterations until the accuracy of the resulting assembly stopped improving or improving slightly. We assessed the improvements using BUSCO scores and the base-level error rate by re-mapping validation long- and short-reads to each assembly (mapped as above). We evaluated the BUSCO scores using BUSCO v3.0.2 with the embryophyta_odb9 lineage (1440 genes in total). Polishing with Racon took between 4 and 12 iterations, and with Pilon between 6 and 10 iterations (Additional file 3: Table S1).

Polishing with both Racon and Pilon significantly improved all of the raw genome assemblies, measured with base-level errors in long- and short-reads, and with BUSCO scores (Additional file 3: Table S1). Polishing with Racon improved long-read base level accuracy by up to 0.83% (in the Marvel_35kb assembly), short-read base level accuracy by up to 1.51% (also in the Marvel_35kb assembly), and the BUSO completeness scores by up to 30.76% (in the Flye_35 assembly). Polishing with Pilon further improved the long-read base level accuracy by up to 0.40% (in the Marvel_35kb assembly), the short-read base level accuracy by up to 1.41% (in the Flye_35kb assembly), and the BUSO completeness scores by up to 24.44% (in the Flye_1kb assembly).

### Assembly ploidy and haplotig removal

Comparison of the polished genome assemblies revealed large variation in assembly size (Table 2). We calculated the assembly ploidy of each assembly as above, assuming a genome size of 500 Mb. The assembly ploidy ranges from 1.12 (Flye_35kb assembly) to 1.79 (Canu_1kb assembly) (Table 2), suggesting that the Canu_1kb assembly is close to a diploid assembly (i.e. ~80% of the genome is represented by two contigs) and that the Flye_35kb assembly is close to a haploid assembly (i.e. only ~12% of the genome is represented by two contigs). To attempt to produce haploid representations of the genome from all assemblies, we used Purge Haplotigs [28] and a custom pipeline, which we call gene conservation informed contig alignment (GCICA) (script available on github from [56]) to find and remove haplotigs from all the assemblies (Fig. 2A).

**Figure 2:**
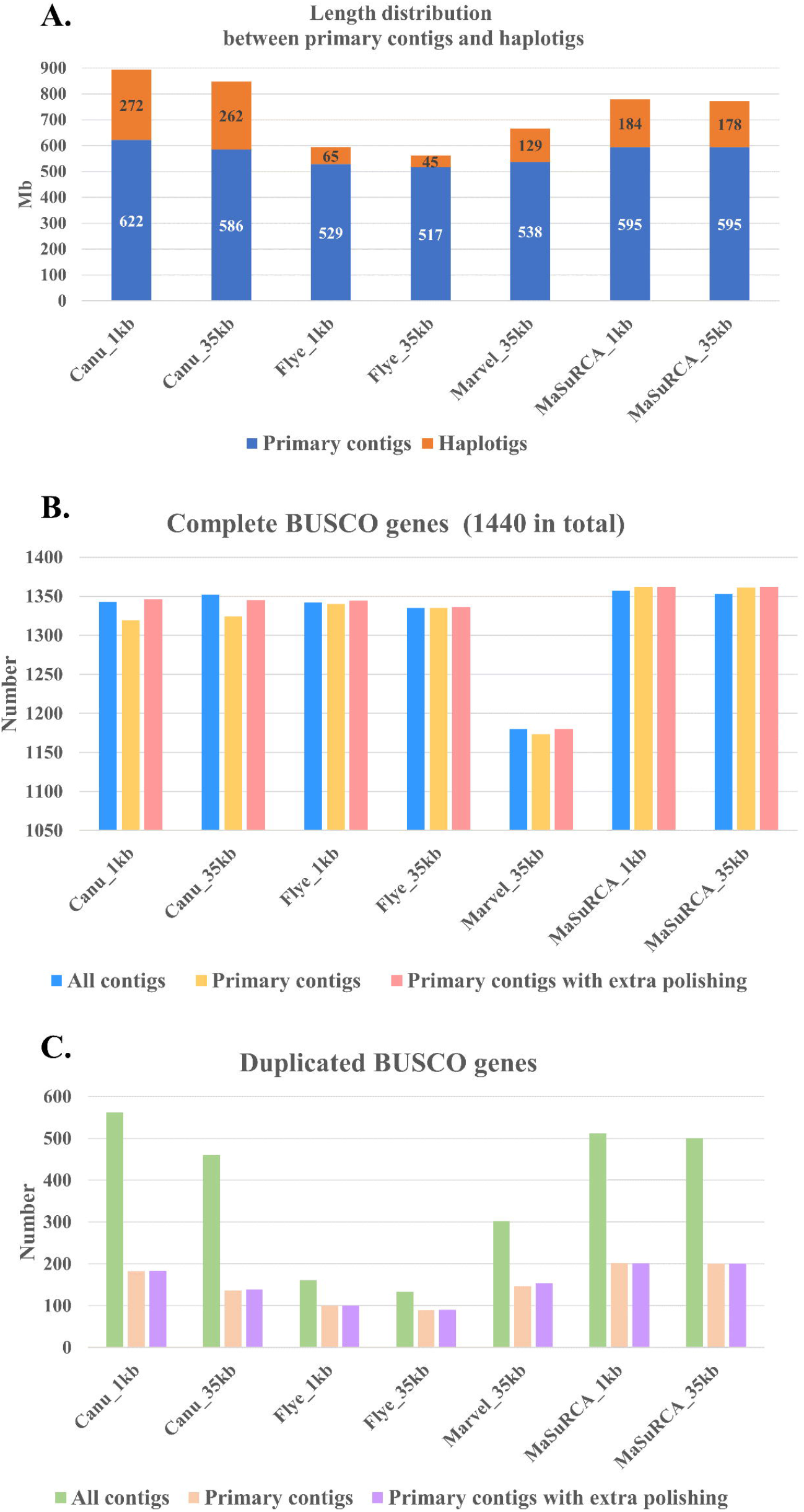
**A.** The length of primary contigs and haplotigs between different assemblies. **B.** The comparison of complete BUSCO genes (1440 in total) between different primary contigs. **C.** The comparison of duplicated BUSCO genes between different primary contigs.

**Figure 3:**
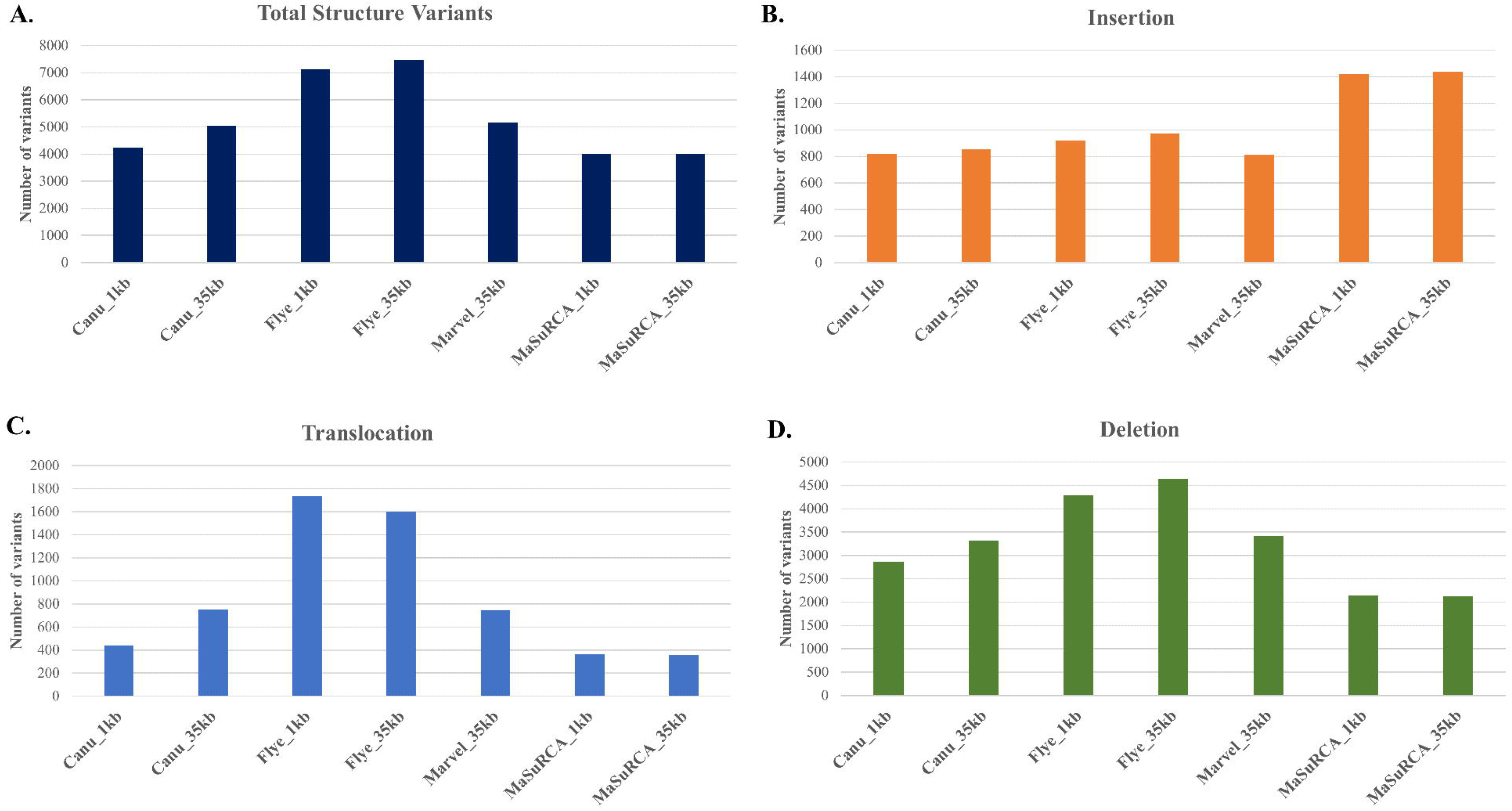
Structural variation analysis of different assembly primary contigs. Each variant was supported by at least 10 long-reads. **A.** The total event of each structural variances of each assembly. **B.** The insertion event of each assembly. **C.** The translocation event of each assembly. **D.** The Deletion event of each assembly.

**Table 2.** Genome size and assembly ploidy

|  | Genome size<br>(bp) | Assembly ploidy | Genome size after<br>Purge Haplotigs (bp) | Assembly ploidy | Genome size after Purge Haplotigs and<br>GCICA (bp)* | Assembly ploidy |
| --- | --- | --- | --- | --- | --- | --- |
| Canu_1kb | 893,781,515 | 1.79 | 645,703,255 | 1.29 | 622,473,836 | 1.24 |
| Canu_35kb | 847,395,928 | 1.69 | 605,520,689 | 1.21 | 586,032,599 | 1.17 |
| Flye_1kb | 593,219,654 | 1.19 | 529,107,244 | 1.06 | 528,619,533 | 1.06 |
| Flye_35kb | 561,597,192 | 1.12 | 517,329,093 | 1.03 | 517,061,277 | 1.03 |
| Marvel_35kb | 666,317,308 | 1.33 | 547,630,224 | 1.10 | 537,813,575 | 1.08 |
| MaSuRCA_1kb | 778,307,850 | 1.56 | 608,764,671 | 1.22 | 594,680,200 | 1.19 |
| MaSuRCA_35kb | 773,071,231 | 1.55 | 608,629,204 | 1.22 | 595,020,257 | 1.19 |

\*Result before final genome polishing.

Purge Haplotigs assigns contigs to primary contigs and haplotigs depending on both coverage information generated by long-read mapping and pairwise alignments of all contigs. To run Purge Haplotigs, we first mapped the long-read assembly dataset to each polished assembly using Ngmlr v0.2.6, and then separated the contigs into primary contigs and haplotigs with default settings. 8% to 29% of each genome assembly was annotated as haplotigs, and removing these haplotigs reduced the assembly ploidy from 1.12 – 1.79 to 1.03 – 1.29 (Table 2).

The high assembly ploidy for some assemblies after running Purge Haplotigs suggested that these assemblies retained haplotigs that covered up to 29% of the genome. We therefore further filtered possible haplotigs using a custom approach, GCICA. If a pair of contigs comprise a primary contig and a haplotig, we would expect most of regions of the haplotig to be very similar to that of the primary contig. To find putative pairs of primary contigs and haplotigs, we therefore looked for pairs of contigs with similar gene content, and then examined these pairs in more detail. To do this, we first mapped the nucleotide sequences of all *E. grandis* genes to all contigs in an assembly using BLASTN v2.7.1+. If >70% of mapped markers in a contig could also be mapped to another contig, and at least 80% of sequence of the smaller contig could be aligned to the other contig (detecting with NUCmer module of MUMmer v4.0.0beta2), we considered these two contigs as a putative primary contig and haplotig pair. We then examined the alignments of all such pairs by eye and removed any pairs in which the smaller contig appeared to be completely contained within the larger, i.e. in which the smaller contig was an unambiguous haplotig. This process identified a further ~2% of each assembly as haplotigs (Table 2).

Following removal of haplotigs, we re-evaluated each assembly using BUSCO scores (Fig. 2). We noted that, depending on the genome assembly, the number of complete BUSCO genes sometimes dropped and sometimes increased slightly after removing haplotigs (Fig. 2B). We hypothesised that BUSCO scores could drop either because haplotig removal mistakenly removed a contig that was not a haplotig, or because haplotig removal correctly removed a haplotig which contained a more conserved representation of a BUSCO gene. BUSCO scores could increase because they are based on E-value scores of alignments, which may be affected by the total length of the assembly. To attempt to alleviate some of these potential issues, we re-polished all of the genome assemblies with multiple rounds of Pilon using the short-read assembly dataset, as above. BUSCO scores recovered across all assemblies with additional Pilon polishing (Fig. 2B). As expected, the number of duplicated BUSCO genes decreased substantially (~50%-70%) after haplotigs were removed from the assemblies and this did not change substantially after additional polishing (Fig. 2C and Additional file 4: Table S2). Together, these results suggest that our haplotig removal pipelines largely succeeded in removing haplotigs, although some haplotigs likely remain if the true genome size is around 500 Mb (Fig. 2A).

### Assessment of assembly quality with eight measures

After haplotig removal and polishing, we considered the primary contigs of each assembly as the final assembly, and evaluated each of the final assembly in using the eight statistics we describe above: contig N50, BUSCO scores, LAI scores, assembly ploidy, base-level error rate, CGAL scores, structural variation and genome sequence similarity (Table 3 and Fig. 4).

**Table 3.** The comparison of final assemblies.

|  |  |  |  | BUSCO score (1440 genes in total) |  |  |  |  |  | LAI<br>scores | Assembly<br>ploidy | Short-read mapping |  | Long-read mapping |  | CGAL<br>scores | Structural<br>variants |
| --- | --- | --- | --- | --- | --- | --- | --- | --- | --- | --- | --- | --- | --- | --- | --- | --- | --- |
|  | Length (bp) | Contig<br>number | Contig<br>N50 (bp) | Complete<br>genes |  | Duplicated<br>genes |  | Fragmented<br>genes |  |  |  | Mapping<br>rate | Error<br>rate | Mapping<br>rate | Error<br>rate |  |  |
| Canu_1kb | 622,218,742 | 895 | 1,502,325 | 1,346 | 93.47% | 183 | 12.71% | 23 | 1.60% | 7.04 | 1.24 | 96.02% | 0.0061 | 91.73% | 0.1661 | -1.959E+06 | 4,243 |
| Canu_35kb | 585,785,283 | 655 | 2,258,674 | 1,345 | 93.40% | 138 | 9.58% | 29 | 2.01% | 5.34 | 1.17 | 95.52% | 0.0066 | 92.64% | 0.1677 | -2.226E+06 | 5,043 |
| Flye_1kb | 528,563,896 | 2,947 | 295,613 | 1,344 | 93.33% | 100 | 6.94% | 31 | 2.15% | 5.7 | 1.06 | 94.86% | 0.0077 | 93.04% | 0.1694 | -2.536E+06 | 7,137 |
| Flye_35kb | 516,992,152 | 2,548 | 385,290 | 1,336 | 92.78% | 90 | 6.25% | 31 | 2.15% | 6.5 | 1.03 | 94.24% | 0.0080 | 92.34% | 0.1699 | -2.726E+06 | 7,458 |
| Marvel_35kb | 537,615,613 | 730 | 1,202,845 | 1,180 | 81.94% | 153 | 10.63% | 32 | 2.22% | 3.77 | 1.08 | 87.37% | 0.0075 | 85.18% | 0.1689 | -4.451E+06 | 5,162 |
| MaSuRCA_1kb | 594,528,099 | 415 | 3,234,447 | 1,362 | 94.58% | 201 | 13.96% | 21 | 1.46% | 9.27 | 1.19 | 94.91% | 0.0060 | 91.57% | 0.1656 | -1.774E+06 | 4,020 |
| MaSuRCA_35kb | 594,871,467 | 416 | 3,234,549 | 1,362 | 94.58% | 200 | 13.89% | 21 | 1.46% | 9.31 | 1.19 | 94.92% | 0.0060 | 91.49% | 0.1655 | -1.790E+06 | 4,017 |

**Figure 4:**
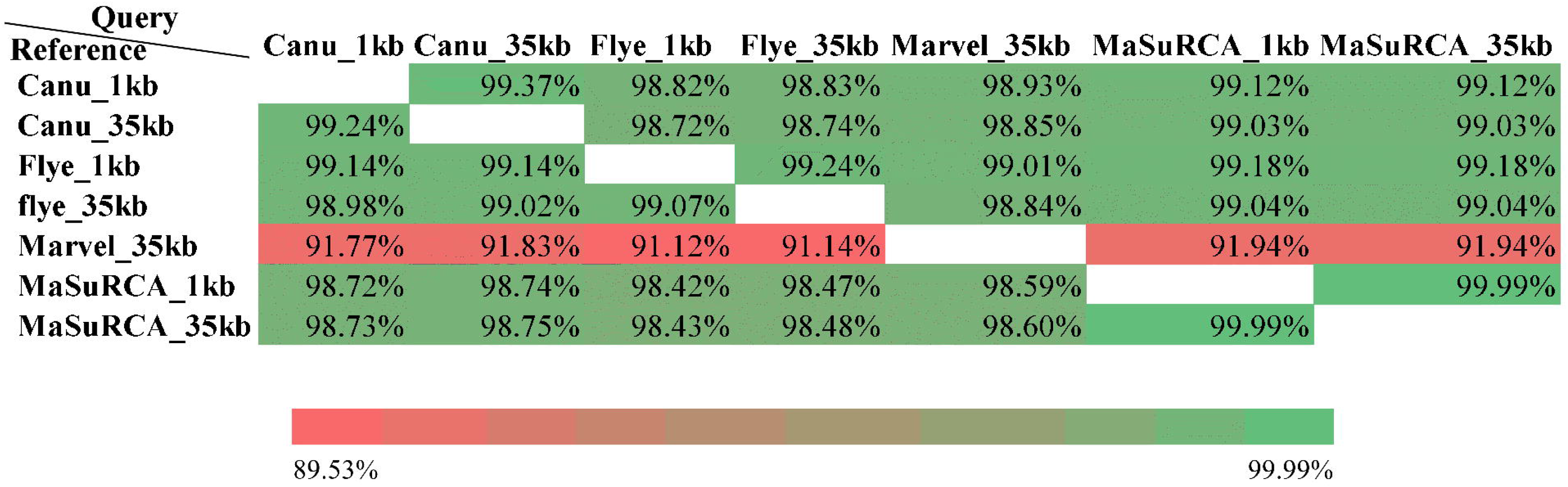
The sequence coverage of whole genome alignment among different assemblies. The sequence coverage was calculated by the length of aligned reference sequence / the total length of reference genome.

Comparison of the eight metrics we used suggested that the MaSuRCA_35kb assembly was likely to be the most accurate assembly overall and that the Marvel_35kb assembly was the least accurate. However, we note that the MaSuRCA assembly did not receive the best scores for all metrics, suggesting that the choice of which assembly to use will sometimes be question-specific. Also, in most of cases, performances of the two MaSuRCA assemblies are very similar.

N50 scores varied from 295 kb (Flye_1kb) to 3.2 Mb (MaSuRCA_35kb), with Flye achieving notably lower N50 values than the other assemblers (Table 3). BUSCO scores ranged from 1180 complete genes (81.94%, Marvel_35kb) to 1362 complete genes (94.58%, MaSuRCA assemblies), although all assemblies except the Marvel_35kb assembly had scores >92%. The MaSuRCA_35kb assembly also achieved the highest LAI score (9.31), which was substantially higher than the best assembly from any other assembler (Canu_1kb, LAI score: 7.04). The lowest LAI score (3.77) was observed in Marvel_35kb assembly. The assembly ploidy was the closest to one for the Flye assemblies (e.g.1.03 for the Flye_35kb assembly vs. 1.19 for the MaSuRCA_35kb assembly). Although these scores have to be interpreted with caution, because the true genome size remains unknown, they are to some extent corroborated by the lower number of duplicated BUSCO genes in the assemblies with the lower assembly ploidy (e.g. 90 duplicated BUSCO genes in the Flye_35kb assembly, vs. 200 in the MaSuRCA_35 assembly). Nevertheless, given that gene duplication is common in *Eucalyptus* species, all such measures need to be interpreted with some caution, since the BUSCO genes themselves could be duplicated in the *E. pauciflora* genome. Taken together, these four metrics suggest that the MaSuRCA_35kb assembly is the most complete, most contiguous, and among the most accurate of the assemblies we produced.

The other three metrics assess the entirety of every assembly, and also suggest that the best assemblies for our data are produced by MaSuRCA (Table 3). The MaSuRCA assemblies (1kb and 35kb) had the lowest error rates (0.006 errors per base for short-read mapping and 0.166 for long-read mapping in both assemblies), and the smallest total number of structural variants estimated from the long validation reads (4017 structural variants for the MaSuRCA_35KB assembly). Flye tended to perform the worst on these metrics, although we note that these results will be affected by the fact that the MaSuRCA assemblies contain more duplicated genome regions (see above), which will tend to reduce the estimated error rates and number of structural variants, because duplicated regions can accurately represent heterozygous variants that will be present in the reads. CGAL ranked MaSuRCA assemblies as the best (1kb: lnL-1774303 and 35kb: lnL-1790386) as the best, and the Marvel_35kb assembly as the worst (lnL-4450742).

Finally, to further investigate the different assemblies, we compared the genome sequence similarity between different assemblies using NUCmer module of MUMmer v4.0.0beta2 (Fig. 4), with the minimum identity set to 75. Notably, around 10% of the sequence of Canu/Flye/MaSuRCA assemblies failed to align to Marvel_35kb assembly (Fig. 4), which, along with the low genome completeness (BUSCO scores) of the Marvel_35kb assembly (Table 3), suggest that the Marvel_35kb assembly may contain many more small duplicated regions than other assemblies. In turn, these duplicated regions may explain the fact that Marvel_35kb assembly has the lowest genome completeness but not the smallest genome size compared to other assemblies (Table 3). Other assemblies have rough 98% - 99% of similarity to each other.

Based on the eight metrics we used above (Table 3), we suggest that the MaSuRCA_35kb assembly represents the most accurate representation of the *E. pauciflora* genome. We note, though, that the Flye assembler only took 1-3% of runtime of the other assemblers used in this paper (Table 1), and produced genome assemblies that were of similar quality to the MaSuRCA_35kb assembly in many respects. The Marvel_35kb assembly received the worst scores on many metrics, and also appears to be missing roughly ~10% of the genome according to BUSCO scores and genome sequence similarity analyses (Table 3).

### Comparative genome analysis between *E. pauciflora* and *E. grandis*

Using the MaSuRCA_35KB assembly, we estimate that the *E. pauciflora* genome is 594,871,467 bp in length, with 416 contigs and a contig N50 of 3,235 kb. The genome has up to 0.006 errors per base. Around 94% of complete BUSCO genes were identified in this *E. pauciflora* genome assembly.

*E. grandis* is the only published *Eucalyptus* genome that is assembled to chromosome level. We therefore compared *E. grandis* with our *E. pauciflora* genome. The *E. grandis* contains 691.43 Mb of sequence, roughly 16% larger than the *E. pauciflora* genome. We compared these two genome assemblies using the NUCmer module of MUMmer v4.0.0beta2 to perform whole genome alignment as described above. This alignment shows that the *E. pauciflora* genome assembly covers just 61.56% of the *E. grandis* genome sequence, leaving approximately 265 Mb of the *E. grandis* genome sequence not covered by the *E. pauciflora* assembly, and 113 Mb of the *E. pauciflora* assembly not covered by the *E. grandis* assembly. Despite this, the coverage of the *E. pauciflora* assembly when mapped to the 11 chromosome-scale scaffolds of the *E. grandis* genome is fairly constant (Fig. 5A), suggesting either that many of these differences result from small errors in both assemblies, and/or from relatively small-scale differences in the underlying genomes.

**Figure 5:**
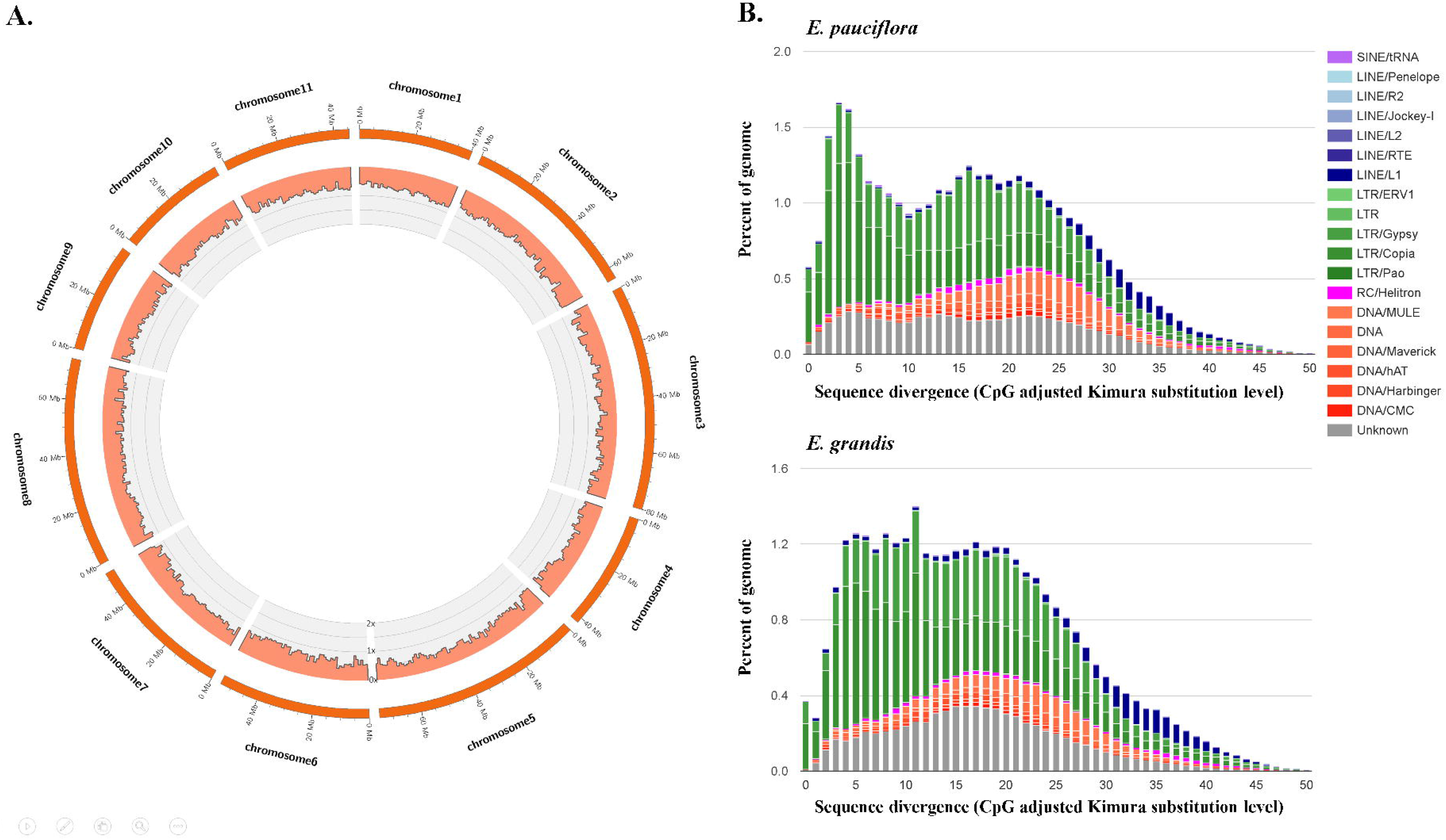
**A.** The histogram of location and coverage of *E. pauciflora* genome aligned to the 11 chromosomes of *E. grandis*. The scale of y-axis is 0x-2x of coverage. Every bar is 1 Mb. The coverage was calculated by the total aligned length of *E. grandis* in each bar / the length of bar. If a site in *E. grandis* is aligned by *E. pauciflora* twice or more, this site will be counted twice or more. **B.** Repeat landscape comparison between *E. pauciflora* and *E. grandis*. Only repeats that are found in both genomes are shown. Older repeat insertions could accumulate more mutations compared to new repeat insertions. This leads to older repeat insertions to have accumulated a higher level of divergence (shown on the right size of the graph).

To examine whether the differences between *E. pauciflora* and *E. grandis* could be explained by their repeat content, we annotated repetitive elements of *E. pauciflora* and *E. grandis* with RepeatMasker v4.0.7 [57]. Although the repeats of *E. grandis* have been annotated before [16], we reannotated them here to enable us to make a direct comparison of the repeat content using an identical pipeline for both genomes. First, we created the custom consensus repeat library using RepeatModeler v1.0.11 [58] with parameter “-engine ncbi”. The classifier was built upon Repbase v20170127 [59]. Then we merged the repeat libraries from RepeatModeler and LTR retrotransposon candidates from LTR retriever to create a comprehensive repeat library as the input for RepeatMasker. We ran the RepeatMasker with “-engine ncbi” model. We used the ‘calcDivergenceFromAlign.pl” script in RepeatMasker pipeline to calculate the Kimura divergence values, and plotted the repeat landscape with repeats presented in both *E. pauciflora* and *E. grandis* genomes.

The repeat content of the two genomes is similar. The *E. pauciflora* genome contains 44.77% of repetitive elements, compared to 41.22% in *E. grandis*. Retrotransposons account for 29.53% of *E. pauciflora* genome, and 26.94% in *E. grandis*, and DNA transposons account for 6.04% and 4.80% of the genome in *E. pauciflora* and *E. grandis*, respectively. The repeat landscapes of the two genomes are also similar, showing roughly two waves of repeat expansion, which is most likely explained by a shared inheritance of most of the repeats in the two genomes (Fig. 5B).

## Conclusions

Here, we report a high-quality draft haploid genome of *E. pauciflora*. It is the first *Eucalyptus* genome assembled with third-generation sequencing reads (Nanopore sequencing), and is the third nuclear genome of *Eucalyptus* species. Due to the economic and ecological importance of *Eucalyptus*, this high-quality genome will support further analysis on *Eucalyptus* and its related species. Additionally, this study will provide useful information for *de novo* plant genome assembly with Nanopore sequencing reads. Finally, the approaches using in this study to assess and compare different assemblies should help in assessing and choosing among many potential genome assemblies

## Availability of supporting data

The *E. pauciflora* genome project was deposited at NCBI under BioProject number PRJNA450887. The whole genome sequencing data are available in the Sequence Read Archive with accession number SRR7153044-SRR7153116. The scripts we used in this paper, including the genome assembly, genome polishing, repeat annotation and genome assessments are available in the Github (https://github.com/asdcid/Eucalyptus-pauciflora-genome-assembly).

## Additional files

**Additional file 1:** A png format with Fig. S1 (GenomeScope result of *E. pauciflora*.)

**Additional file 2:** A png format with Fig. S2 (Genome contamination detection. Almost all sequences were matched the sequences in streptophyta phylum group. No contamination was found.)

**Additional file3:** A xlsx format with Table S1 (The comparison of polishing results of raw assemblies.)

**Additional file4:** A xlsx format with Table S2 (The comparison of polishing result of each genome after haplotig removal.)

## Abbreviations

BUSCO: Benchmarking Universal Single-Copy Orthologs
CGAL: computing genome assembly likelihoods
*E. grandis*: *Eucalyptus grandis*
*E. pauciflora*: *Eucalyptus pauciflora*
NCBI: the National Center for Biotechnology Information
LTR: long-terminal repeat
LAI: long-terminal repeat assembly index

## Ethics Statement

*E. pauciflora* leaves were collected a single *E. pauciflora* individual in Thredbo, Kosciuszko National Park, New South Wales, Australia (Latitude –□ 36.49433, Longitude 148.282983). The written permission was from the Scientific Licensing office of the Office of Environment and Heritage for New South Wales: www.licence.nsw.gov.au, in accordance with national guidelines in Australia. Tissues were not deposited as voucher specimens.

## Author Contributions

AD, DK, RL and WW conceived this project. AMS and RL performed sample collection for Illumina sequencing. AMS extracted genomic DNA, and constructed library for Illumina sequencing. RL and MS carried out sample collection for Nanopore sequencing. MS and BS performed DNA extraction, library preparation, and Nanopore sequencing. DK performed long-read polishing and Canu 1kb assembly, whereas AD performed Canu_35kb, Flye_1kb Flye_35kb and Marvel_35kb assemblies and contamination detection. AD and WW conducted the whole genome alignment analysis. WW conducted all the remaining analyses. AD, BS, DK, RL and WW were involved in data interpretation. AD, RL and WW drafted the original manuscript. RL and WW finalized the manuscript. All authors read and approved the final manuscript.

