## Supplementary figures and images for "The draft nuclear genome assembly of *Eucalyptus pauciflora*: new approaches to comparing *de novo* assemblies"

### Fig. S1

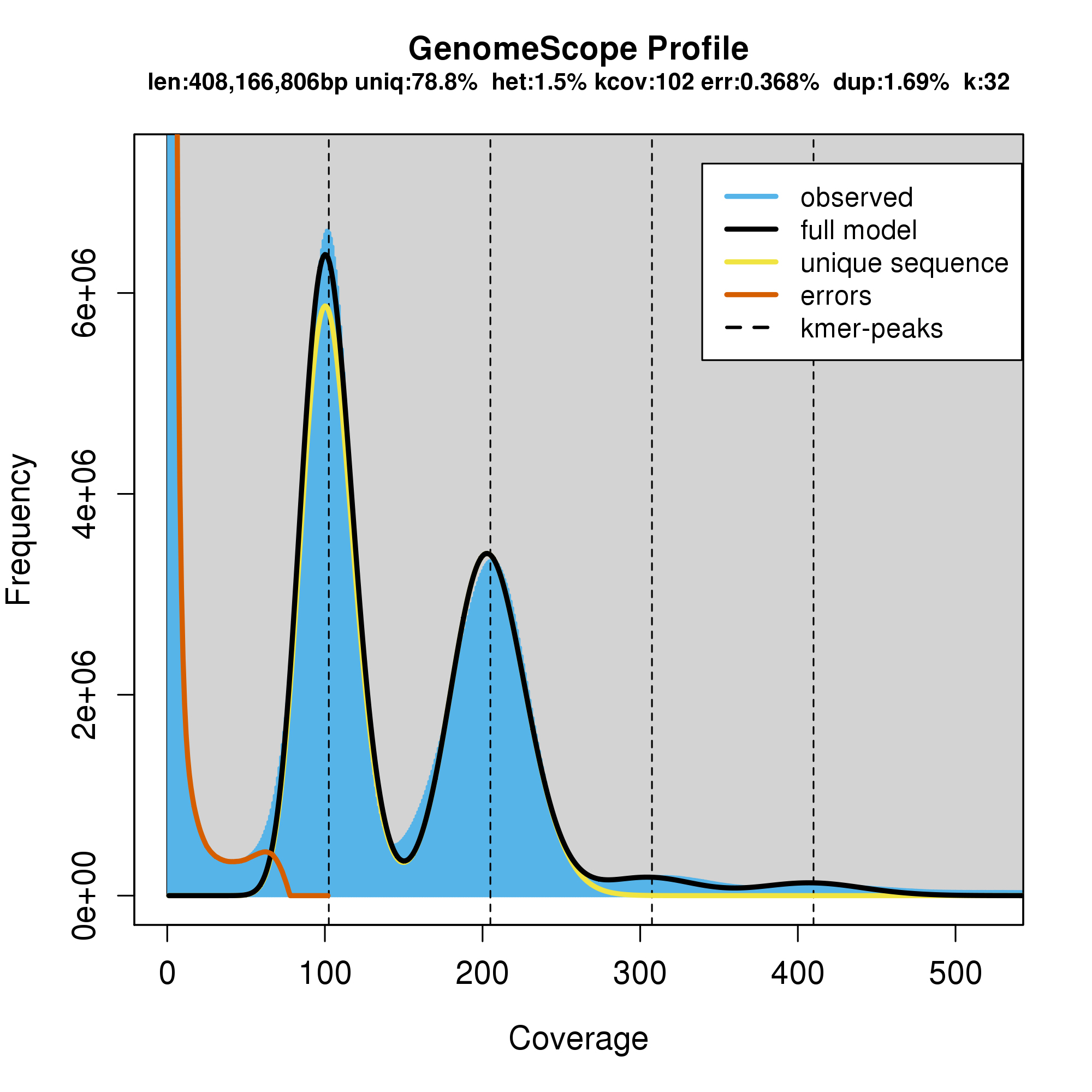

### Fig. S2

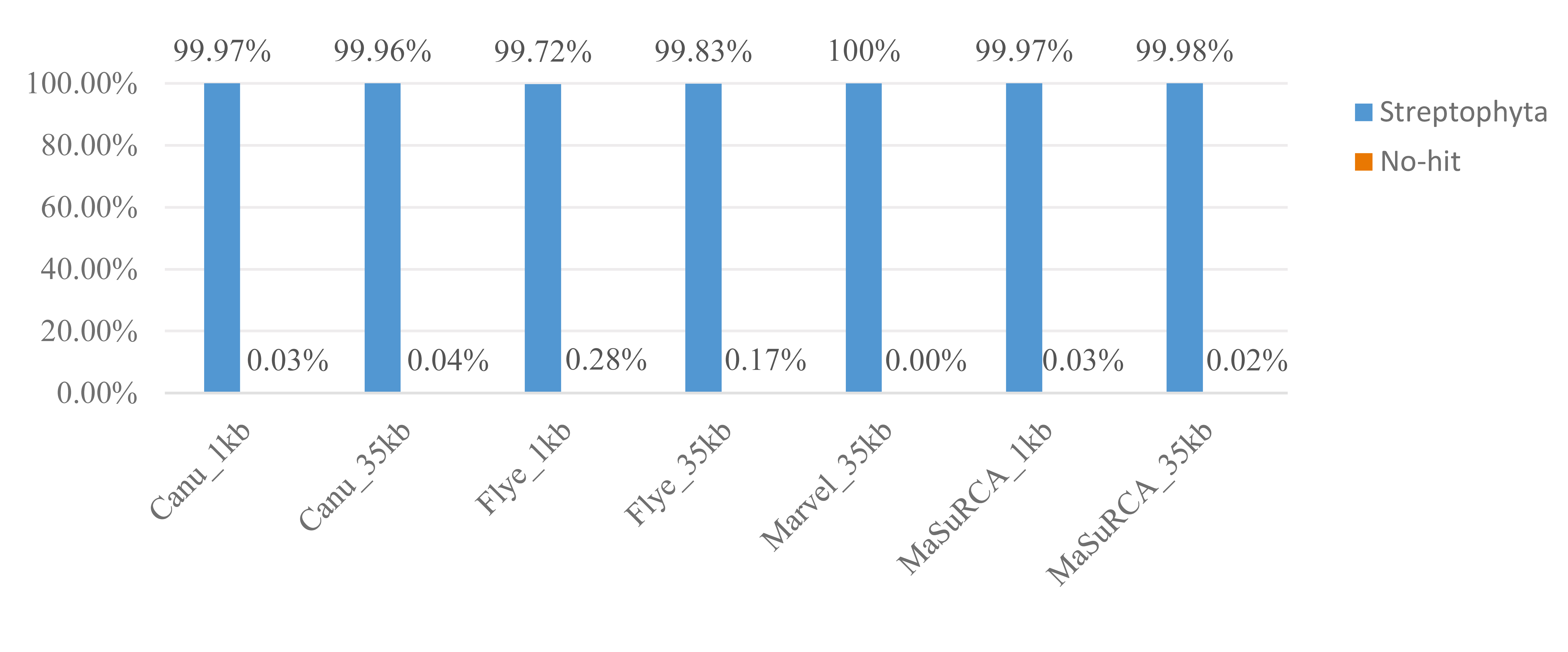
